# Enrichment of novel *Verrucomicrobia, Bacteroidetes* and *Krumholzibacteria* in an oxygen-limited, methane- and iron-fed bioreactor inoculated with Bothnian Sea sediments

**DOI:** 10.1101/2020.09.22.307553

**Authors:** Paula Dalcin Martins, Anniek de Jong, Wytze K. Lenstra, Niels A. G. M. van Helmond, Caroline P. Slomp, Mike S. M. Jetten, Cornelia U. Welte, Olivia Rasigraf

## Abstract

Microbial methane oxidation is a major biofilter preventing larger emissions of this powerful greenhouse gas from marine coastal areas into the atmosphere. In these zones, various electron acceptors such as sulfate, metal oxides, nitrate or oxygen can be utilized. However, the key microbial players and mechanisms of methane oxidation are poorly understood. In this study, we inoculated a bioreactor with methane- and iron-rich sediments from the Bothnian Sea in order to investigate microbial methane and iron cycling under low oxygen concentrations. Using metagenomics, we observed shifts in the microbial community over approximately 2.5 years of bioreactor operation. Marker genes for methane and iron cycling, as well as respiratory and fermentative metabolism, were investigated. Metagenome-assembled genomes representing novel *Verrucomicrobia, Bacteroidetes* and *Krumholzibacteria* were recovered and revealed potential for methane oxidation, organic matter degradation, and iron cycling, respectively. This work brings new insights into the identity and metabolic versatility of microorganisms that may be members of such functional guilds in coastal marine sediments and highlights that the methane biofilter in these sediments may be more diverse than previously appreciated.

**Importance:** Despite the essential role of microorganisms in preventing most methane in the ocean floor to reach the atmosphere, comprehensive knowledge on the identity and the mechanisms employed by these microorganisms is still lacking. This is problematic because such information is needed to understand how the ecosystem functions in the present and how microorganisms may respond to climate change in the future. Here, we enriched and identified novel taxa potentially involved in methane and iron cycling in an oxygen-limited bioreactor inoculated with methane- and iron-rich coastal sediments. Metagenomic analyses provided hypotheses about the mechanisms they may employ, such as the use of oxygen at very low concentrations. The implication of our results is that in more shallow sediments, where oxygen-limited conditions are present, the methane biofilter is potentially composed of novel, metabolically versatile *Verrucomicrobia* that could contribute to mitigating methane emissions from coastal marine zones.

## Introduction

Archaea and bacteria capable of methane oxidation largely prevent global emissions of methane, a greenhouse gas 28-105 times more potent than carbon dioxide for global warming, into the atmosphere (1). In deep marine sediments, archaeal methanotrophs are predicted to consume more than 90% of the *in-situ* generated methane in cooperation with sulfate-reducing bacteria (2). In this way, a biofilter that prevents larger methane emissions from these ecosystems is established. Recent estimates suggest that 45-61 Tg of methane are oxidized annually in marine sediments with sulfate, the dominant terminal electron acceptor for methane oxidation in such environments (3). However, methanotrophs operating under low oxygen concentrations and using alternative electron acceptors such as iron and manganese oxides, nitrate or even limited amounts of oxygen are poorly identified, and the mechanisms and metabolism they employ are not yet well explored. Characterizing these microorganisms and understanding their environmental functioning is fundamental to estimate impacts of climate change and eutrophication in coastal sediments, to design more predictive models, and to create possible future bioremediation and restoration strategies.

Iron oxides are globally distributed in marine costal sediments (4). Anaerobic oxidation of methane coupled to iron reduction (Fe-AOM) is hypothesized to account for elevated dissolved iron concentrations in methanic zones, particularly in the Baltic and Bothnian Sea, North Sea and Black Sea (4). However, despite strong biogeochemical evidence for Fe-AOM (5–7), this remains one of the least elucidated methane-cycling metabolisms. Archaea affiliated to the cluster ANME-2d, of the genus *Candidatus* Methanoperedens, were the first identified microorganisms to show methane oxidation activity at the expense of Fe and manganese (Mn) reduction, likely reversing the methanogenesis pathway (8). *Candidatus* Methanoperedens nitroreducens, the first archaeon described to couple methane oxidation to nitrate reduction (9, 10), was shown to perform Fe- and Mn-AOM in short-term experiments with iron citrate, nanoparticulate ferrihydrite and birnessite in follow-up studies (8). Recent investigations (11, 12) enriched related Fe- and Mn-AOM Methanoperedens species, namely *Ca*. Methanoperedens ferrireducens, *Ca*. Methanoperedens manganicus, and *Ca*. Methanoperedens manganireducens from organic-rich freshwater sediments in Australia after two years of cultivation. The genomes of the various *Ca*. Methanoperendes strains encode several multiheme *c*-type cyctochrome proteins that are implicated in the extracellular electron transfer pathways needed to convey electrons to the metal oxides (12–15).

Interestingly, bacteria commonly implicated in aerobic methane oxidation via particulate methane monooxygenase (PMO) have very recently been suggested to be capable of Fe-AOM via a yet unknown mechanism. Pure cultures of the gammaproteobacterial and alphaproteobacterial methanotrophs *Methylomonas* and *Methylosinus* were shown to couple methane oxidation to ferrihydrite reduction under the availability of 0.89 mg O_2_ L^-1^ (16). Bacterial methanotrophs were suggested to account for Fe-AOM in oxygen-depleted incubations with sediments from Lake Kinneret in Israel (17) and in anoxic waters of Northwestern Siberian lakes (18). How methane may be activated by PMO in the absence of oxygen or at nanomolar concentrations of oxygen is not yet known, but could have similarities to the mechanism employed by *Ca*. Methylomirabilis species (19, 20). How pure cultures of these methanotrophs could reduce iron while their genomes lack known marker genes for iron reduction is another question that remains unanswered.

The brackish Bothnian Sea is located in the northern part of the Baltic Sea and, in contrast to the rest of the Baltic Sea basin, is an oligotrophic ecosystem. These conditions have established due to the topography, which hinders input of nutrient-rich waters from the south. The Bothnian Sea is fed by several major rivers that transport freshwater, terrestrial organic carbon and metal oxides into the ecosystem (21). Low salinity and high sedimentation rates have enabled the establishment of a relatively shallow sulfate-methane transition zone (SMTZ). Below the SMTZ, ferric oxides (ferrihydrite) can accumulate and act as terminal electron acceptors for AOM (6). A recent study from another area in the Baltic Sea, Pojo Bay estuary in Finland, also provided evidence for AOM below the SMTZ in Fe-rich coastal sediments (22). The microbial communities from both ecosystems exhibited significant similarities, including dominant taxa such as ANME-2a/b, *Methanomicrobia, Bathyarchaeota, Thermoplasmata, Bacteroidetes, Chloroflexi, Verrucomicrobia*, and *Proteobacteria* (α-, β-, γ-) (22, 23). However, a direct link between particular taxa and Fe-AOM activity in Baltic Sea sediments is still lacking.

Previous metagenomic and biogeochemical studies have indicated that a variety of electron acceptors and different guilds of microorganisms, including various putative methanotrophs, are present in Bothnian Sea sediments (6, 23, 24) and could play a role in carbon, sulfur, nitrogen, and iron cycling. To better understand the metabolism and ecophysiology of these organisms, we inoculated a bioreactor with oxygen-depleted sediment from an iron and methane-rich sediment from a coastal site in the Bothnian Sea. Previous incubation experiments (6) and modelling studies (25, 26) have indicated Fe-AOM activity in these sediments. The reactor received methane and ferric iron (both Fe(III) nitrilotriacetic acid and ferrihydrite nanoparticles) in order to enrich microorganisms involved in methane oxidation and iron reduction. Microbial community dynamics were followed with metagenomics for approximately 2.5 years. This cultivation effort resulted in the enrichment of novel *Verrucomicrobia, Bacteroidetes* and *Krumholzibacteria*, which are hypothesized to be involved in methane oxidation, organic matter degradation and iron cycling, respectively.

## Methods

### Bioreactor setup

The enrichment culture was operated in sequencing batch mode in a jacketed 3L-glass bioreactor (Applikon, Delft, The Netherlands) at a working volume of 2 L. Medium (0.3 L day^-1^) was continuously supplied, except during daily settling (1h) and effluent pumping out times (30 minutes). The reactor was inoculated on 20 June 2016 with 411 g of wet sediment collected on 6 August 2015 from sampling site NB8 in the Bothnian Sea (N 63°29.071, E 19°49.824) (26). The sediment was derived from 37-42 cm depth, located below the SMTZ, where the pore water sulfate concentration was below the detection limit (<75 µM), but where methane and Fe(II) concentrations were in the millimolar range, and iron oxides were abundant (26). Between sampling and inoculation, the sediment was stored anoxically at 4°C in the dark in a sealed aluminum bag under dinitrogen gas pressure. The medium consisted of 0.1 mM KH_2_PO_4_, 2 mM KCl, 3 mM CaCl_2_, 80 mM NaCl, 9.5 mM MgCl_2_, 0.2 mM NH_4_Cl, 5 mM Fe(III) nitrilotriacetic acid (NTA), and the trace element solution (2 mL / 10 L) was made as previously described (27) and supplemented with 0.2 mM Ce_2_(SO_4_)_3_. The Fe(III)NTA solution (200 mM) was made according to the following protocol: 57.3 g of NTA was added to 200 mL MilliQ water and the pH was adjusted to 7-8 with 10M NaOH until the NTA was dissolved. After addition of 16.4 g NaHCO_3_ and 27.0 g FeCl_3_x6H_2_O to the dissolved NTA, the volume was adjusted to 500 mL with MilliQ water. The solution was made sterile by passing it through a 0.2 μm filter. Additionally, the reactor received 10-12 mM of Fe(OH)_3_ (ferrihydrite) nanoparticles, synthesized as previously described (28), once a month from 7 Aug 2017.

The medium was constantly sparged with an Ar/CO_2_ gas mixture (95:5) and the culture was continuously sparged with a CH_4_/CO_2_ gas mixture (95:5, the Linde Group, The Netherlands) with a flow rate of 10 mL min^-1^. The liquid volume was maintained by a level-controlled effluent pump, the stirring was set at 150 r.p.m., and the reactor was kept at room temperature (21°C). The pH was monitored using an ADI 1010 Bio Controller (Applikon, Delft, The Netherlands) and maintained at pH 7.59 by a pH controller loop using potassium hydrogen carbonate (KHCO_3_) as base and CO_2_ gas as acid. Oxygen was monitored by a Clark-type oxygen electrode (Applikon, Delft, The Netherlands) and measured during activity assays as described below. During the standard operation mode of the bioreactor, oxygen concentrations were below the detection limit of the electrode. To prevent growth of photosynthetic organisms and to prevent the reduction of iron by UV light, the reactor was wrapped in black foil, and black tubing with low oxygen permeability was used (Masterflex Norprene, Cole Parmer, USA).

### Activity assays

Whole reactor activity tests were conducted twice, one in 2016 and one in 2017, as follows. Medium supply, effluent outflow, and base pump were stopped and the reactor was flushed with Ar/CO_2_ (95:5) while stirring. Methane concentrations were measured in the headspace, and when undetectable (below 1.8 ppm), ^13^CH_4_ was added to a final concentration of 10% in the headspace. Fe(III)NTA (5-10 mM) or Fe(OH)_3_ nanoparticles (10 mM) were tested as terminal electron acceptors. For batch activity tests, which were performed in 2018, 12.8-15 mL of reactor biomass was placed into 30 mL-serum bottles and incubated with a combination of electron acceptors and donors (in duplicates). The following conditions were set up: all bottles were kept at 0.5 bar overpressure, electron donors were ^13^CH_4_ at 75% of the headspace or 2 mM acetate, and electron acceptors were 15 mM Fe(OH)_3_ nanoparticles, 15 mM Fe(III) citrate, 2 mM magnesium sulfate, or O_2_ at 5% of the headspace. Control bottles included biomass and only methane or oxygen.

Headspace samples (100 µL) were withdrawn with a gas tight glass syringe (Hamilton, Switzerland) and methane was immediately measured in technical triplicates on a HP 5890 gas chromatograph equipped with a Porapak Q column (80/100 mesh) and flame ionization detector (Hewlett Packard, Palo Alto, CA, USA). For ^13^CO_2_ and O_2_ technical duplicate measurements, 50 µL of headspace was injected into an Agilent 6890 series gas chromatograph coupled to a mass spectrometer (Agilent, Santa Clara, CA, USA) equipped with a Porapak Q column heated at 80°C with helium as the carrier gas. Gas concentrations were calculated using a calibration curve made with gas standards, and liquid-dissolved concentrations were estimated with the Ostwald coefficient (29). Iron concentrations were measured in technical duplicates using the colorimetric ferrozine method (30). Briefly, 30 μL of a solid-free reactor liquid sample was mixed with 30 μL 1M HCl in an eppendorf tube (for ferrous iron), and in another tube 30 μL of solid-free liquid sample was mixed with 30 μL of saturated hydroxyl amine solution in 1M HCl (for total iron). After incubation of 1 hour at room temperature, 10 μL of the solution was added to 100 μL ferrozine reagents (0.05% wt. / vol. ferrozine (PDT disulfonate: 3-[2-Pyridyl]-5,6-diphenyl-1,2,4-trizaine-4,4-disulfonic acid sodium salt) in 50 mM 4-(2-hydroxyethyl)-1-piperazineethanesulfonic acid (HEPES) buffer, pH 7.0) and 500 μL of MilliQ water. The absorbance was measured at 562 nm using a SpectraMax 190 microplate reader (Molecular Devices, Sunnyvale, CA, USA). The ferric iron content was calculated by subtracting the concentration of ferrous iron from the concentration of the total reduced ferrous iron. Activity data was imported into RStudio v1.2.5033 (R v3.6.3) (31) and graphs were constructed with ggplot2 v3.3.2 (32).

### DNA extraction, metagenomic sequencing, data processing, and analyses

DNA was extracted from original sediments (herein referred to as T0) and from biomass after 16 months (T1) and 29 months (T2) of reactor operation. All DNA extractions were performed using the DNeasy PowerSoil DNA extraction kit (Qiagen, Hilden, Germany) according to the manufacturer’s instructions. One extra DNA extraction of T2 with the ammonium acetate method (33) was conducted and used only for assembly and binning purposes. Metagenomic sequencing was performed in-house using the Illumina Nextera® XT Library Prep Kit according to the manufacturer’s instructions (Illumina, San Diego, CA, USA). The library was normalized to 4 nM and sequencing was performed with an Illumina MiSeq using the sequencing protocol for 300 bp paired-end reads. Resulting reads (∼4 Gbp per sample) were trimmed and quality-controlled with BBDuk (https://sourceforge.net/projects/bbmap/), then co-assembled with MEGAHIT v1.2.9 (34). Read mapping was performed with BBMap (https://sourceforge.net/projects/bbmap/) and mapping files were handled with samtools v1.9 (using htslib v1.9) (35). Contigs were binned with four methods all using default parameters: binsanity v2.0.0 (36), concoct v1.1.0 (37), maxbin2 v2.2.7 (38) and metabat2 v2.15 (39). Bins were supplied to DAS tool v1.1.2 (40) for consensus binning and to CheckM v1.1.2 (41) for quality inference of metagenome-assembled genomes (MAGs).

Taxonomic classification of MAGs was obtained with GTDB-Tk v0.3.2 (42), and phylogenetic trees for MAG placement were constructed using UBGC v3.0 (43). MAGs were gene-called with prodigal v2.6.3 (44) and annotated with KEGG KAAS (45), prokka v1.13.3 (46), and DRAM v0.0.1 (47) in KBase (48). Marker genes for iron metabolism were searched with FeGenie (49), while other genes of interest were searched using hmmsearch (HMMER 3.3 with --cut_tc) (50), blastp (51), prokka and DRAM annotation files. The following hmms were downloaded from PFAM (https://pfam.xfam.org/) or TIGRFAMs (http://tigrfams.jcvi.org/cgi-bin/index.cgi): PF02240.16 (MCR_gamma), PF02241.18 (MCR_beta), PF02783.15 (MCR_beta_N), PF02249.17 (MCR_alpha), PF02745.15 (MCR_alpha_N), PF14100.6 (PmoA), PF04744.12 (PmoB and AmoB), PF02461.16 (AMO), TIGR04550 (MmoD), PF02406.17 (MmoB/DmpM family), and PF02964.16 (Methane monooxygenase, hydrolase gamma chain). Average amino acid identity was calculated using the Kostas lab enveomics tool (52) available at http://enve-omics.ce.gatech.edu/aai/index.

Phylogenetic trees for specific genes of interest were constructed by retrieving reference sequences from NCBI (https://www.ncbi.nlm.nih.gov/protein/), aligning them with MUSCLE v3.8.31 (53), stripping alignment columns with trimAl v1.4.rev22 (54) (with -gappyout) and calculating trees with FastTree v2.1.10 (55). Trees were visualized on iTol (56). Reads per kilobase per million mapped reads (RPKM) values for genes of interest were calculated by mapping reads to these genes with BBMap using the option rpkm. Data frames were imported into RStudio v1.2.5033 (R v3.6.3) (31) and heat maps were constructed using the packages vegan v2.5-6 and ggplot2 v3.3.2 (32), with the function heatmap.2. All figures were edited in Adobe Illustrator CC 2018 (Adobe, San Jose, California, USA).

### Data availability

Raw sequencing reads, metagenome-assembled genomes, and unbinned contigs have been deposited on NCBI under the BioProject accession number PRJNA663545.

## Results and Discussion

### Retrieved metagenome-assembled genomes revealed microbial community shifts over reactor operation time

Over the course of 29 months, the occurrence of iron reduction was obvious from the color change in the reactor and production of iron (II) in activity tests (Figure S1). Consumption of methane especially at the expense of iron (III) was less apparent (Figure S1). We did observe production of ^13^CO_2_ from ^13^CH_4_, but could not ascertain that this was coupled to stoichiometric iron reduction. Most surprisingly, after more than 2 years of operation under low headspace oxygen concentrations, generally between 70 and 0.4 µM (calculated liquid-dissolved oxygen between 4.7 and 0.27 µM), we still detected quite high levels of oxygen-dependent methane oxidation (Figure S1).

In order to investigate the changes in the microbial community, original sediments and bioreactor samples were subjected to DNA extractions and metagenomic sequencing. Illumina sequencing, co-assembly, and binning allowed the reconstruction of 56 MAGs (Figure 1) from three time points: original sediments (T0), bioreactor biomass 16 months after reactor inoculation (T1), and 29 months after reactor inoculation (T2). Together, these 56 MAGs represented 35.8%, 81.5%, and 79.1% of metagenome reads at T0, T1, and T2, respectively. In original sediments, three MAGs represented 6-9% of metagenome reads each: thermoplasmata_1, aminicenantales_1, and bipolaricaulia_1. Using percent of reads mapped to each genome as a proxy for abundance, these organisms seemed to disappear by T2 (Table S1). At T1, 31% of metagenome reads mapped to the MAG desulfuromonadales_1, potentially representing the most abundant organism in the reactor at that time, followed by bacteroidales_2, with 8.6%, and burkholderiales_1, with 6.9%. By T2, these microorganisms appeared to become rare members of the community. Finally, at T2, three MAGs accounted for 29.7% of metagenome reads: the verrucomicrobial MAG pedosphaeraceae_1, with 10.5% of reads, bacteroidales_8 with 10.1%, and krumholzibacteria_1 with 9.1% (Table S1).

**Figure 1.**
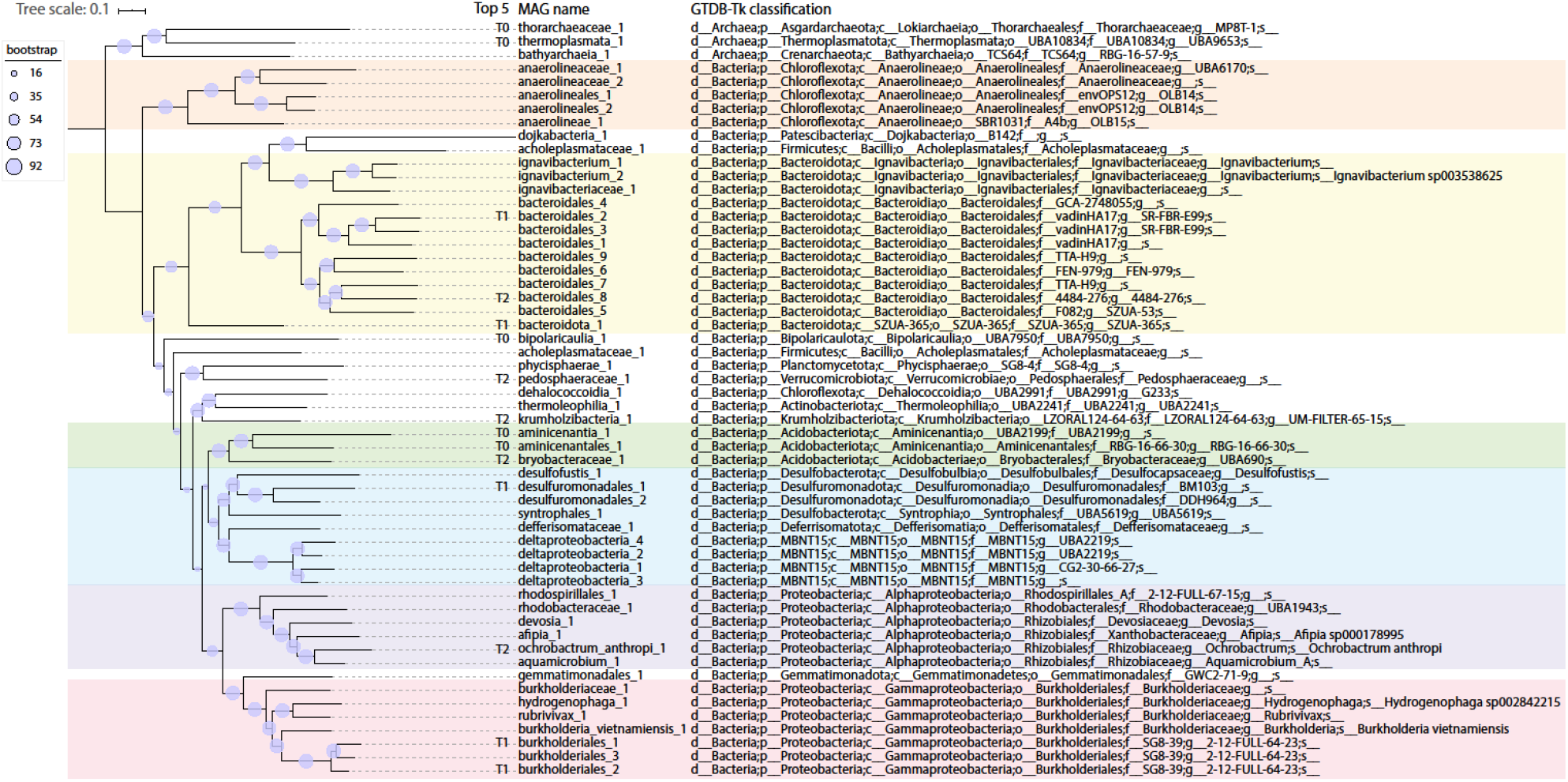
UBCG phylogenetic tree of MAGs retrieved in this study. Ninety-two conserved genes were used for tree construction (43). MAGs are named after their taxonomy, which was assigned with GTDB-Tk (42). The top five most abundant MAGs in each metagenome (T0, T1 and T2) are indicated. Colors highlight the following taxa: orange, *Anaerolineae*; yellow, *Bacteroidota*; green, *Acidobacteriota*; blue, *Deltaproteobacteria*; purple, *Alphaproteobacteria*; pink, *Gammaproteobacteria*.

The MAGs retrieved in this study generally reflect findings from previous metagenomic and 16S rRNA gene analyses of sediments from site NB8 (23), which were used as inoculating material for the bioreactor in this study. *Anaerolinea* and *Bacteroidetes* were detected at increasing 16S rRNA gene-based relative abundances with depth in NB8 and were hypothesized to perform fermentation. Other groups previously identified in NB8 via 16S rRNA gene analyses (23) included *Verrucomicrobia, Planctomycetes, Ignavibacteria, Actinobacteria, Alphaproteobacteria* (particularly *Rhodospirillales* and *Rhizobiales*), *Deltaproteobacteria* and *Gammaproteobacteria*. A previously recovered MAG classified as *Aminicenantes* had potential for acetogenesis, sulfate reduction and fermentation, while *Thorarchaeota* and *Bathyarchaeota* MAGs were hypothesized to participate in fermentative production of acetate, formate and ethanol (23). Similarly, other studies (24, 57–59) that sequenced DNA from Baltic Sea water or sediments have identified several common groups, including *Acidobacteria, Actinobacteria, Bacteroidetes, Chloroflexi, Firmicutes, Gemmatimonadetes, Planctomycetes, Alphaproteobacteria, Deltaproteobacteria, Gammaproteobacteria, Thermoplasmata* and *Verrucomicrobia*.

Shifts in microbial community composition detected in this study via several metagenomic analyses (further detailed in the next sections) highlight microbial successions over time and might be explained by a combination of factors. Firstly, we hypothesize that, due to high terrestrial influence in the Bothnian Sea, sediments used as inoculum carried recalcitrant organic matter (60) that could have been gradually degraded, providing differing pools of carbon at times and thus affecting microbial community structure. Secondly, carbon fixation and biomass turnover might also have provided different pools of organic matter into the reactor. Finally, monthly shots of Fe(OH)_3_ nanoparticles in addition to the constant inflow of Fe(III)NTA starting approximately one year after reactor inoculation may have changed the bioreactor conditions periodically.

### Marker gene analyses highlight potential for methane and iron cycling, as well as oxygen respiration

Genes of interest were searched in MAGs in order to investigate functional potential for methane and iron cycling, as well as respiratory metabolisms (oxygen reduction, denitrification, and sulfate reduction) and carbon fixation pathways. The analyzed marker genes for methane cycling were subunits of the methyl-coenzyme A reductase (*mcrA, mcrB*, and *mcrG*), soluble methane monooxygenase (*mmoB* and *mmoX*) and particulate methane monooxygenase (*pmoA* and *pmoB*). RPKM values derived from read mapping to genes were used as proxy for gene abundance across the three metagenomes.

No canonical soluble (*mmo*) or particulate (*pmo*) methane monooxygenase genes were found in this study’s entire dataset. Five *mcr* genes with best blastp hits to the ANME-2 cluster archaeon assembly GCA_009649835.1 were identified in unbinned contigs (three in the same one). RPKM values indicate the organism represented by these genes was abundant in original sediments but was selected against in the reactor (Figure S2). While it is unclear why archaea did not thrive in the reactor, we hypothesize that keeping the reactor at atmospheric pressure and room temperature (21°C) might have played a role as methane dissolves less in the liquid phase in comparison to a pressurized setting at colder temperatures. The identification of *mcr* genes aligns with previous 16S rRNA gene analyses of NB8 sediments (23), which revealed ANME-2a/b dominated archaeal communities at depths 14-30, 34-45 and 50-63 cm in this site (in the SMTZ and below as well). Archaea affiliating to this group were hypothesized to mediate SO_4_^2-^- and Fe-AOM in distinct sediment zones (23), directly reducing iron or, alternatively, transferring electrons to an iron-reducing organism.

Next, we looked at C1 metabolism. Genes encoding nicotinamide adenine dinucleotide (NAD)-dependent methanol dehydrogenases were found in the following 13 MAGs: bryobacteraceae_1, thermoleophilia_1, aminicenantales_1, burkholderiales_1, burkholderiales_2, burkholderiales_3, anaerolineaceae_1, deltaproteobacteria_1, deltaproteobacteria_2, desulfofustis_1, devosia_1, rhodospirillales_1, and pedosphaeraceae_1. Shifts in microbial community were also apparent from RPKM values of methanol dehydrogenases over time (Figure S3). By T2, MAG pedosphaeraceae_1 had the methanol dehydrogenase with the highest RPKM value, followed by rhodospirillales_1, and deltaproteobacteria_1.

Marker genes for iron oxidation and iron reduction were searched with FeGenie (49). Marker genes for iron oxidation identified in some MAGs of this study (Figure 2) encoded the following: *cyc2*, iron oxidases; *cyc1*, periplasmic cytochrome *c*_4_, part of an iron oxidation complex; *foxE*, a *c*-type cytochrome; and *foxY*, a protein containing the redox cofactor pyrroloquinoline quinone (PQQ). Other MAGs had marker genes for iron reduction that encoded several porins and outer membrane *c*-type cytochromes described in a variety of iron-reducing microorganisms (49). RPKM values for these genes highlighted microbial community shifts over time in the reactor (Figure 2). The desulfuromonadales_1 MAG appeared to be particularly abundant in T1, when the reactor started to receive monthly inputs of Fe(OH)_3_ nanoparticles in addition to the constant inflow of Fe(III)NTA provided in the medium. At T2, when the supply of ferrihydrite was stopped, the functional guild of iron reduction seemed to be spread across other members represented by MAGs deltaproteobacteria_4, ignavibacteriaceae_1, deltaproteobacteria_1, and krumholzibacteria_1. As for iron oxidation, three MAGs had *cyc2* genes: aminicenantales_1 and aminicenantia_1, which may represent iron-oxidizing microorganisms in original sediments, and krumholzibacteria_1, which seemed abundant at the latest enrichment culture metagenome (T2). These results are supported by whole reactor activity tests performed 6 months after reactor inoculation, which provided evidence for iron reduction (Figure S1). While we did not conduct iron oxidation activity tests, we hypothesize iron (II) produced from iron (III) reduction could have fueled iron oxidation in the bioreactor. Traces of oxygen could have been the possible electron acceptor for this process. While *Deltaproteobacteria* and *Ignavibacteria* are known iron reducers (61, 62), *Aminicenantes* have only recently been suggested to perform iron oxidation due to the identification of *cyc2* gene sequences in draft genomes (49), corroborating findings from this study. Given the widespread occurrence of *Aminicenantes* bacteria in Bothnian Sea sediments (23), their contribution to iron cycling could be significant. The potential for iron reduction (via OmcS) and oxidation (via Cyc2) in *Krumholzibacteria* is described here for the first time.

**Figure 2.**
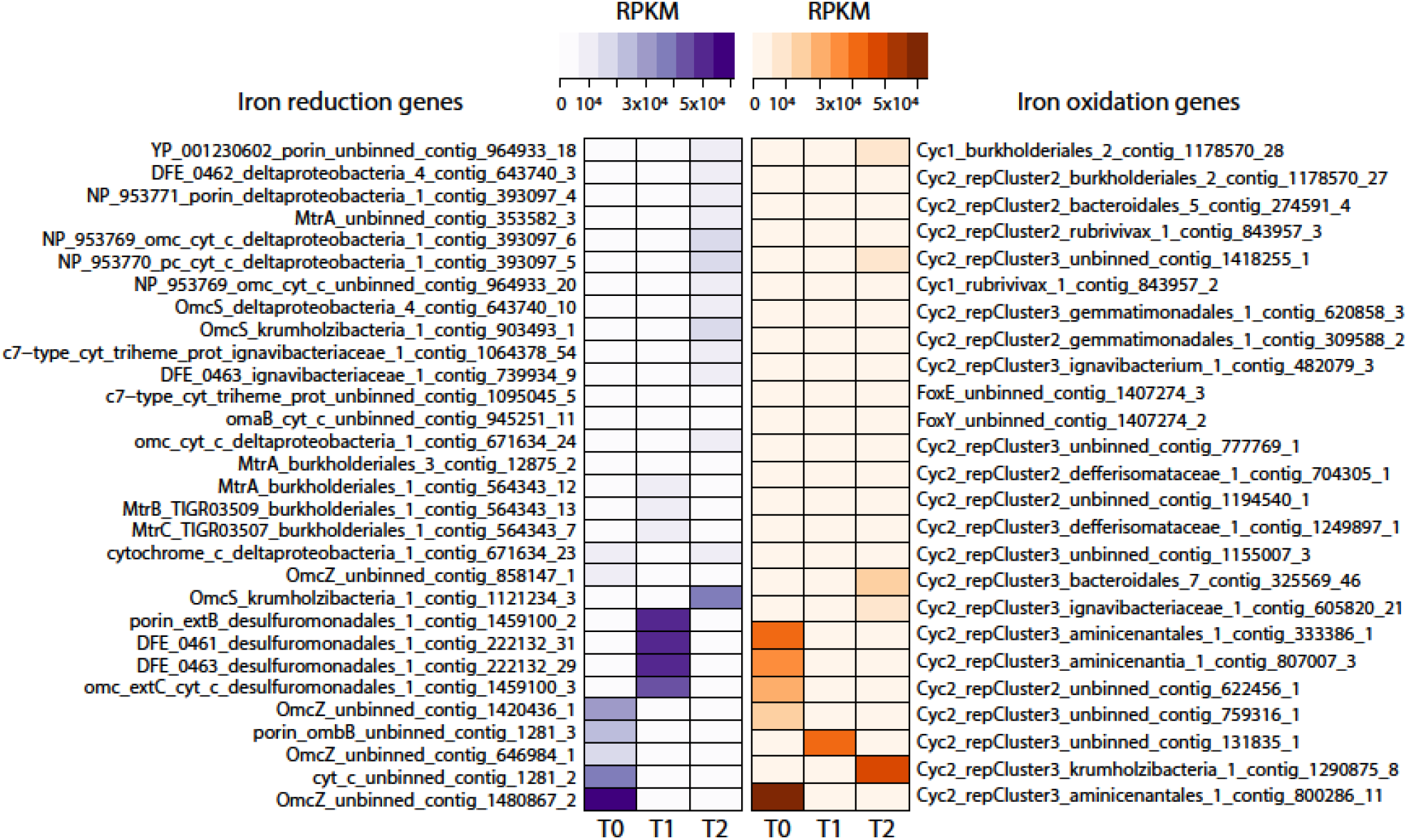
Heat maps of marker genes for iron reduction and iron oxidation in the three metagenomes (T0, T1, and T2). Marker genes were identified with FeGenie (49). RPKM values were calculated from read mapping with BBMap (https://sourceforge.net/projects/bbmap) and imported into RStudio (31) for heat map construction.

MAGs were mined for potential utilization of terminal electron acceptors other than iron (Table S1). The investigated marker genes encode enzymes involved in denitrification (*nar, nir, nor, nos* and *nrf*), sulfate reduction (*dsr*) and oxygen respiration (several quinol and cytochrome *c* oxidases). The gene *dsrA* was found in 8 MAGs, although in desulfofustis_1 it likely encodes a protein subunit involved in sulfur oxidation. Marker genes for denitrification and/or dissimilatory nitrate reduction to ammonium (DNRA) were present in 48 MAGs, but none had the full denitrification pathway. Potential for oxygen respiration was also widespread: 37 MAGs had *cox* genes, 19 had *cbb*_*3*_-type subunit-encoding genes, and 32 had *cyd* genes. The widespread functional potential for the utilization of alternative terminal electron acceptors has been previously reported: potential for dissimilatory sulfate reduction was detected in previous NB8 MAGs affiliated to *Bacteroidales, Xanthomonadales*/*Chromatiales, Aminicenantes, Syntrophobacterales*, and *Gemmatimonadales* (23). In an investigation of Bothnian Sea site US5B, *narG, napA, nirK, nirS, nor, nosZ*, and *nrfA* were identified across the depth profile, but seemed particularly abundant at shallower sediments, where nitrogen oxides would be available for respiration (24).

The persistence of oxygen respiration potential in oxygen-depleted sediments may occur upon rapid burial of metabolically versatile microorganisms due to high sedimentation rates, as reported for our study site (26). In surface sediments where oxygen is still available (here: 0-3 mm depth in summer (63)), oxygen respiration is expected to be the predominant metabolic mode. As microorganisms are buried and oxygen becomes depleted, alternative electron acceptors and metabolic pathways might then be employed by these microorganisms.

Genes involved in carbon fixation pathways were investigated in order to determine potential for autotrophy, which could have accounted for inputs of organic matter into the reactor (Table S1). Our results indicate widespread potential for carbon fixation among MAGs: ribulose bisphosphate carboxylase (RuBisCO) genes were used as markers for the Calvin-Benson-Bassham cycle and were identified in 7 MAGs. Genes encoding 2-oxoglutarate synthase and pyruvate-ferredoxin/flavodoxin oxidoreductase, markers for the reverse tricarboxylic acid (TCA) pathway, were both found in 30 MAGs. Genes for acetyl-CoA/propionyl-CoA carboxylase, markers of the 3-hydroxypropionate cycle, were found in 22 MAGs. Finally, the carbon monoxide dehydrogenase/acetyl-CoA synthase, marker for the reductive acetyl-CoA pathway, was encoded in 8 MAGs.

### Enrichment of a novel *Verrucomicrobia, Bacteroidetes*, and *Krumholzibacteria* microorganisms

The MAG pedosphaeraceae_1 likely represented an abundant microorganism enriched after ∼2.5 years of reactor operation. The estimated genome completeness was 98.6%, with 8.8% redundancy. This MAG had most genes encoding enzymes in the aerobic pathway for methane oxidation, with the exception of a canonical methane monooxygenase. However, pedosphaeraceae_1 harbored 7 *pmoA*-family sequences that, in a phylogenetic tree with verrucomicrobial reference sequences, formed a cluster (indicated by the number 2, green highlight, in Figure 3) with other hypothetical proteins and *pmoA*-family sequences. Another cluster (indicated by 1, purple highlight, in Figure 3), monophyletic with cluster 2, contained mostly canonical reference *pmoA* sequences, while sequences not in these clusters were mostly reference *pmoB*.

**Figure 3.**
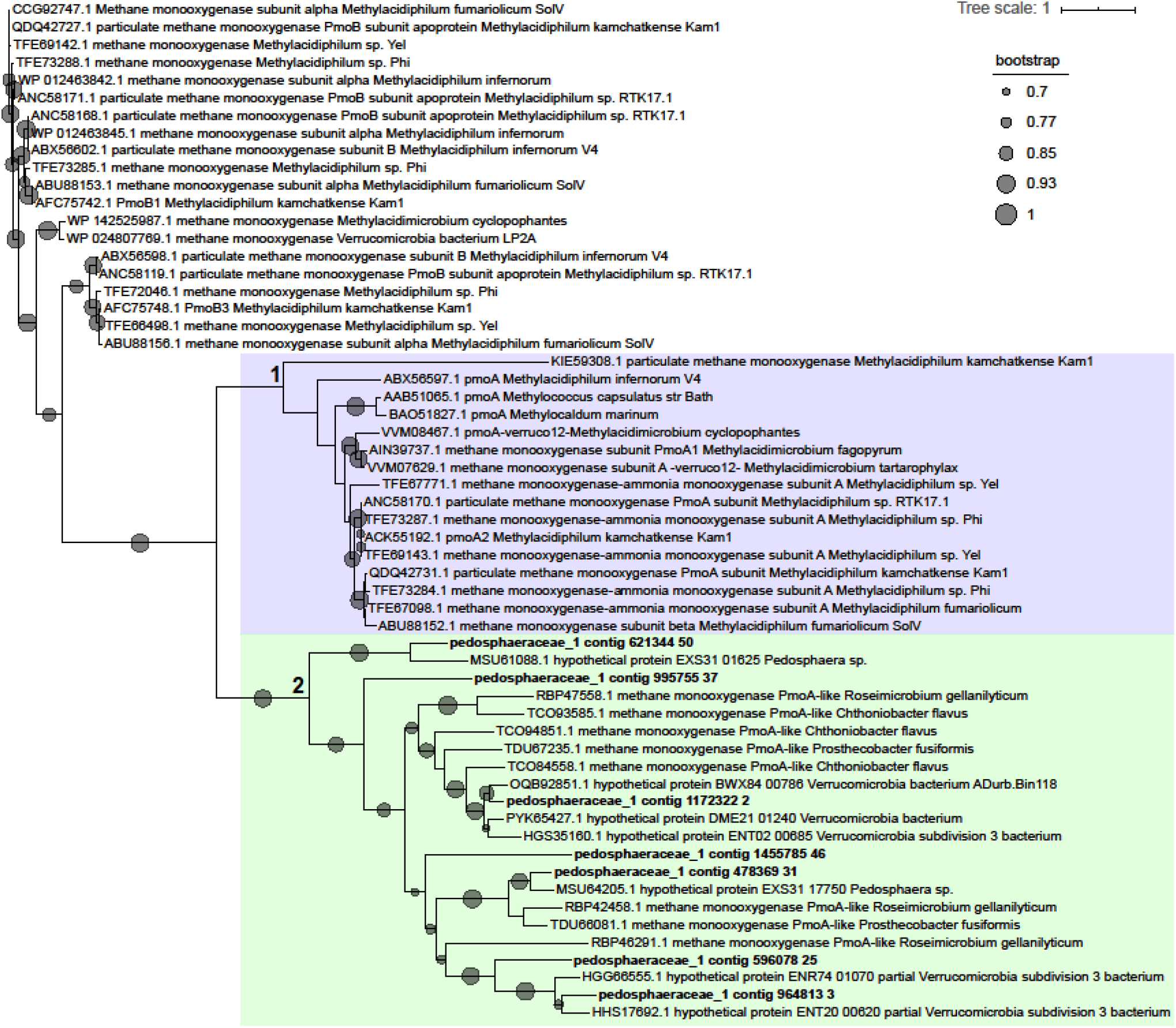
Phylogenetic tree of PmoA-family sequences found in the MAG pedosphaeraceae_1 and reference verrucomirobial sequences. Reference sequences were retrieved from the NCBI RefSeq database and are indicated by accession numbers. Sequences were aligned with MUSCLE (53), alignment columns were stripped with trimal (54), and the tree was built with FastTree (55) using the Jones-Taylor-Thornton model of amino acid evolution (64).

Other genes in the pathway for aerobic methane oxidation included an NAD-dependent methanol dehydrogenase, the three proteins involved in the tetrahydrofolate pathway for formaldehyde oxidation to formate, and a formate dehydrogenase gamma subunit (other subunits were missing). Electron transport chain proteins for oxygen respiration were mostly present: all subunits of the NADH dehydrogenase (type I), subunits *sdhABC* encoding succinate dehydrogenase, oxygen reductase-encoding genes *cyoE, ctaA, coxC, cbb*_*3*_-type subunits I/II and III, and *cydAB*. Nitrogen dissimilatory metabolism genes *narGH, napAB, norB*, and *nrfAH* were also present in the genome, as well as all genes in glycolysis, the TCA cycle and the pentose phosphate pathway. Marker genes for the reverse TCA cycle (*korAB, por/nifJ, porB*) and for the 3-hydroxypropionate cycle (*accABCD*), as well as the *acs* gene encoding acetyl-CoA synthetase, indicated potential for carbon fixation. The metabolic potential of pedosphaeraceae_1 is summarized in Figure 4.

**Figure 4.**
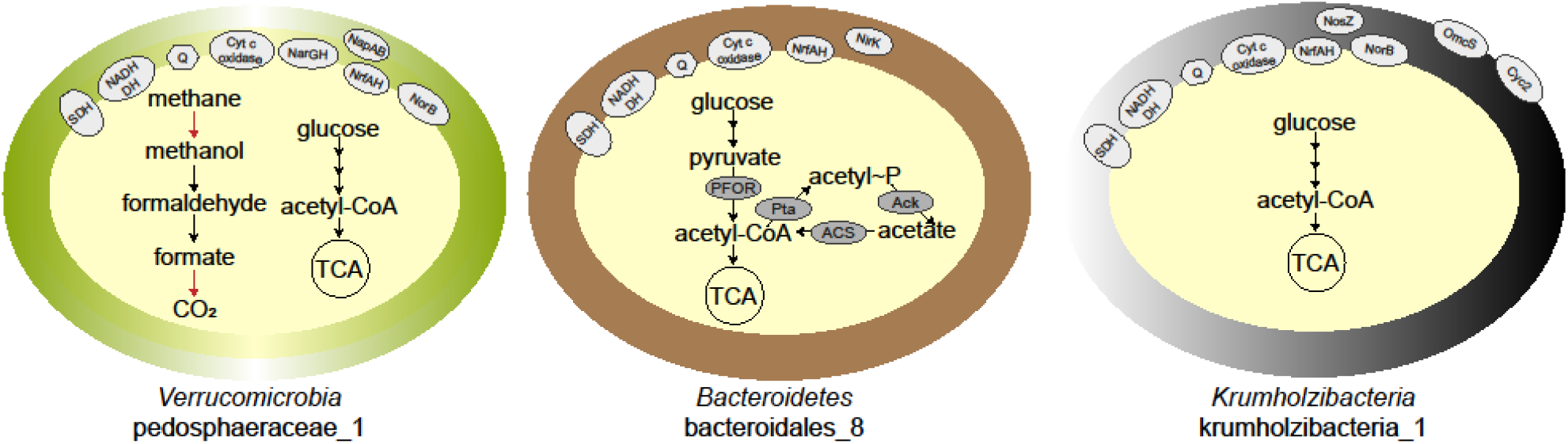
Highlights of metabolic potential found in the three enriched microorganisms after ∼2.5 years of bioreactor operation. MAG name and taxonomy are indicated below each cartoon. Red arrows indicate the protein was not clearly identified or subunits are missing. Abbreviations are as follows: SDH, succinate dehydrogenase; NADH DH, NADH dehydrogenase; Q, quinone; TCA, tricarboxylic acid cycle; PFOR, pyruvate:ferredoxin oxidoreductase; Pta, phosphotransacetylase; Ack, acetate kinase; ACS, acetyl-CoA synthetase, Nar, nitrate reductase; Nap, periplasmic nitrate reductase, Nir, nitric oxide-forming nitrite reductase; Nor, nitric oxide reductase; Nos, nitrous oxide reductase; Nrf, ammonia-forming nitrite reductase; OmcS, outer membrane *c*-type cytochrome; Cyc2, iron oxidase.

By T2 (29 months after reactor inoculation), when pedosphaeraceae_1 was most abundant, *mcr* genes present in unbinned contigs had low RPKM values (∼2000-2500) in comparison to *pmoA*-family genes present in pedosphaeraceae_1 (∼15000-23000). Activity tests suggest that the bioreactor biomass was actively oxidizing methane 6 months after reactor inoculation, and that the biomass was responsive to oxygen, which promoted methane oxidation in serum bottle incubations performed in the second year of reactor operation (Figure S1). These results support the role of a methane-oxidizing microorganism thriving under the low dissolved oxygen concentrations (4.7 - 0.27 µM) generally measured in the reactor (Figure S1). Although activity data did not coincide with DNA extraction times, the presence of high affinity oxygen reductases in the genome indicated that MAG pedosphaeraceae_1 could have been responsible for methane oxidation potentially coupled to oxygen reduction. We hypothesize that a divergent particulate methane monooxygenase in the PMO family or a canonical methane monooxygenase missing from the genome could have accounted for the activity of methane oxidation. Methanol leakage from pedosphaeraceae_1 could have sustained other community members such as bryobacteraceae_1 and rhodospirillales_1, which also had methanol dehydrogenases and represented 4.3% and 3.5% of metagenome reads at T2, respectively. Alternatively, pedosphaeraceae_1 might have thrived on methanol oxidation, which leaves the questions of what was the source of methanol and which microorganism oxidized methane in the bioreactor.

A previous investigation (23) showed that verrucomicrobial 16S rRNA gene sequences dominated bacterial communities at site NB8 (and at the nearby site N10) in the top 4 cm of the sediment, but were also present throughout the entire depth profile of NB8 at lower abundances. In this same top 4cm-depth, few sequences of canonical methane-oxidizing proteobacteria were identified (23). *Verrucomicrobia* were suggested to degrade polysaccharides and algal material while performing aerobic metabolism or denitrification at shallower depths. In deeper sediments, however, oxygen and sulfate are depleted, and iron reduction is predicted to predominate (26). This indicates that *Verrucomicrobia* may be metabolically versatile and capable of surviving under oxygen-depleted conditions, and that the enrichment of such microorganisms has relevance for biogeochemical cycling *in-situ*. We hypothesize that *Verrucomicrobia* potentially oxidizing methane with a distinct methane monooxygenase might thrive in a niche in shallow sediments where the transition from low to zero oxygen occurs (Figure 5). In deeper sediments, where oxygen is depleted and the redox potential is low, archaea would dominate methane oxidation (23). Our results indicate that the methane biofilter in these coastal marine sediments could be more complex than previously appreciated, comprising also novel methanotrophs within the *Verrucomicrobia* adapted to low oxygen availability.

**Figure 5:**
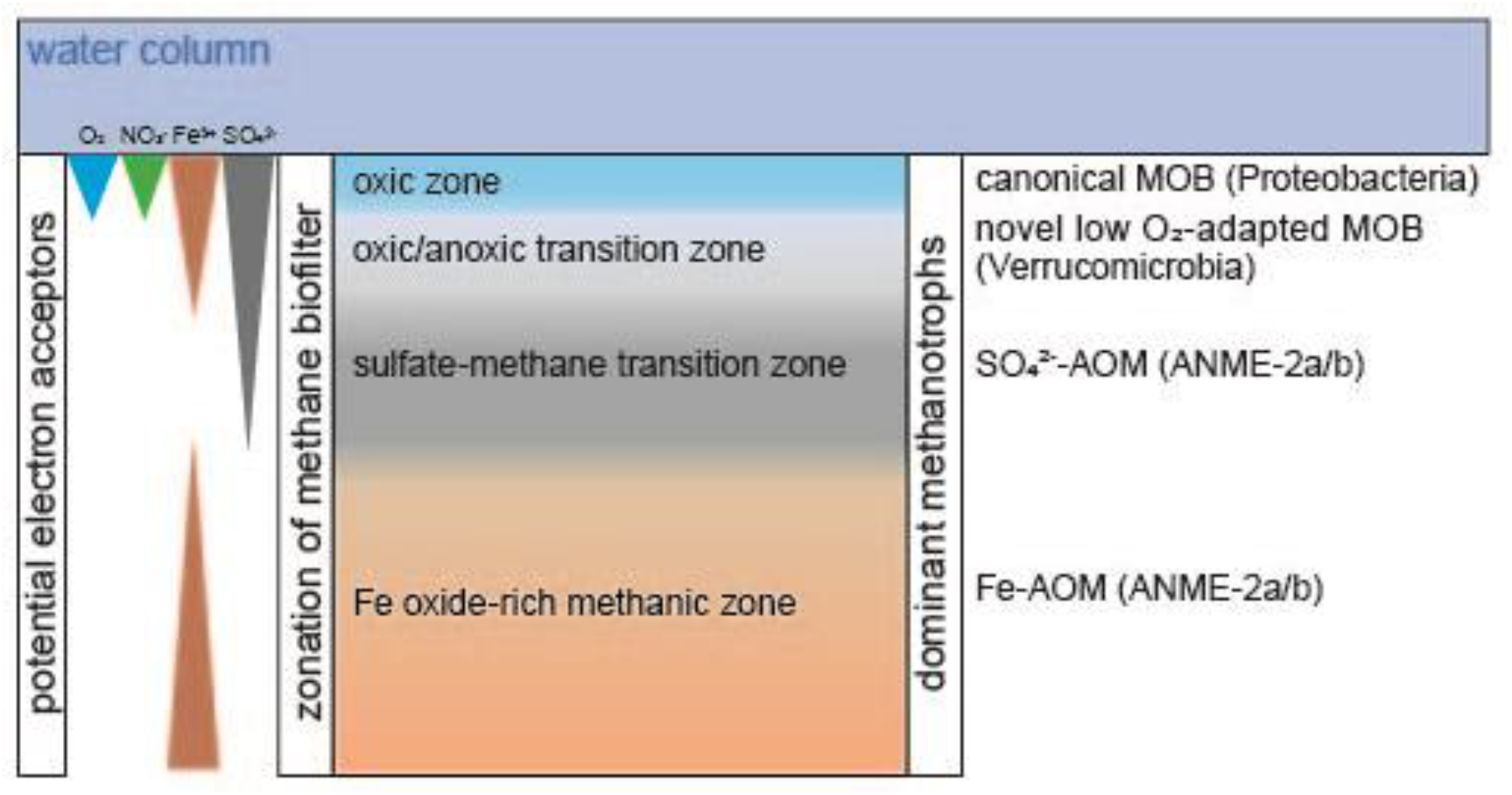
Illustration of the hypothesized biological methane filter in the coastal sediments of the Bothnian Sea. The depth of distinct zones and electron acceptor availability is not to scale. Abbreviations: MOB, methane-oxidizing bacteria; ANME, anaerobic methane-oxidizing archaea; AOM, anaerobic oxidation of methane.

An important question remains regarding whether pedosphaeraceae_1 and, more generally, bacteria utilizing particulate methane monooxygenase for methane oxidation could couple it to iron reduction. Under such scenario, very low concentrations of oxygen could suffice for methane activation by PMO. If pedosphaeraceae_1 performed Fe-AOM in the bioreactor, the produced Fe(II) could have served as electron donor for *Krumholzibacteria*. This highly speculative hypothesis is theoretically based on a recent study, which demonstrated that two proteobacterial methanotrophs reduced ferrihydrite during growth on methane under low oxygen concentrations, in the absence of known marker genes for iron reduction in both genomes (16). Evidence for this phenomenon became first available from Lake Kinneret sediment incubations with ^13^C-methane and different iron oxide minerals under oxygen depletion (17). In that study, *pmoA* copy numbers followed trends of δ^13^C_DIC_ and iron (II) indicative of Fe-AOM. More recently, *Methylomonadaceae* were suggested to account for methane oxidation in anoxic waters of Northwestern Siberian lakes (18). The authors hypothesized that iron (III) could be the electron acceptor for AOM because of total iron concentrations and the identification of taxa known to perform iron reduction. Adding to that, nitrate was below their detection limit and nitrite concentrations did not exceed 0.02 mg L^-1^ of N-NO_2_^-^ which indicated the unavailability of these electron acceptors or their rapid turnover.

While the family *Pedosphaeraceae* has no isolates or genomes previously implicated in methane oxidation, several known methanotrophs within the *Verrucomicrobia* phylum have been reported, such as *Methylacidiphilum fumariolicum* SolV (65) and others of the same genus, as well as some species of *Methylacidimicrobium* (66). Our study highlights the metabolic versatility of organisms in the phylum and indicates *Verrucomicrobia* might play unforeseen roles in marine environments. Further investigations are needed to elucidate the metabolism of these organisms.

MAG bacteroidales_8 (96.2% complete, 1.6% redundant) had potential for oxygen respiration via low and high affinity oxygen reductases-encoding genes (*coxBCD, cbb*_*3*_-type subunits I/II, III, and IV, *and cydAB*), as well as DNRA via NrfAH (Table S1), and nitrite reduction to nitric oxide via NirK (Figure 4). Marker genes for the reverse TCA cycle (*korAB, por/nifJ*) were present. As most genes in the glycolysis and TCA pathways were found, we hypothesize that this organism could have been involved in heterotrophic respiratory metabolism. *Bacteroidetes* have been previously identified in water and sediment samples from the Baltic and Bothnian Sea (23, 24, 59). *Bacteroidetes* previously detected in NB8 were hypothesized to perform fermentation, which is also supported by the presence of a phosphate acetyltransferase, acetate kinase, acetyl-CoA synthetase, pyruvate:ferredoxin oxidoreductase-encoding gene *pfor*, and the NADP-reducing hydrogenase HndABCD in bacteroidales_8. In another study, nine *Bacteroidetes* MAGs retrieved from water samples of the Baltic Sea also had cytochrome *c* oxidase genes (data retrieved from annotations on NCBI) and were hypothesized to contribute to the degradation of algal material via carbohydrate active enzymes (58), which has similarly been reported for *Bacteroidetes* from the North Sea (67). Given that, as expected in the oligotrophic NB8 site, no cyanobacteria were detected in our metagenomes, it is more likely that the organism represented by MAG bacteroidales_8 could have participated in biomass turnover evidenced by the observed shifts in MAG abundances over time in the bioreactor. Whether respiratory or fermentative metabolism was employed by this microorganism remains to be elucidated.

MAG krumholzibacteria_1 (89% complete, 2.5% redundant) seemed to be involved in iron cycling, harboring OmcS and Cyc2-encoding genes that had high RPKM values by T2 (Figures 2 and 4). It is intriguing that krumholzibacteria_1 had both iron oxidation and reduction genes. The capacity for iron oxidation and reduction in the same organism has been previously described in *Geobacter sulfurreducens* (68, 69). Further studies are necessary to elucidate these potential roles of krumholzibacteria_1 in iron cycling. If krumholzibacteria_1 coupled iron oxidation to oxygen reduction, it must have been able to compete for the limited oxygen present in the bioreactor. Respiratory metabolism genes present in this MAG indicated this could have been possible: we identified genes encoding subunits of low and high affinity oxygen reductases (*cox, cbb*_*3*_-type, *cyd*, and *cyo*). Additionally, *norB, nosZ*, and *nrfAH* were present. Arsenic resistance genes *arsC, arsB, arsM* and *arsR* were present, as well as complete glycolysis and most genes in the TCA pathway. Potential for carbon fixation was determined based on the presence of marker genes for the reverse TCA cycle (*korAB, porABC*) and for the 3-hydroxypropionate cycle (*accABCD*), as well as the *acs* gene encoding acetyl-CoA synthetase. The genome also had a hydroxylamine dehydrogenase *hao* gene.

The functional potential of krumholzibacteria_1 differs from that of *Candidatus* Krumholzibacterium zodletonense Zgenome0171^T^ (QTKG01.1), type material for the recently described phylum *Krumholzibacteriota*, recovered from the sulfur-rich Zodletone spring in southwestern Oklahoma, USA (70). For comparative purposes in this study, the draft genome QTKG01.1 has been annotated with DRAM (47). While krumholzibacteria_1 presented potential to respire iron, oxygen, nitric oxide, nitrous oxide, and nitrite, QTKG01.1 showed no clear potential to utilize inorganic external terminal electron acceptors. However, QTKG01.1 harbored a few subunits of the NADH:quinone oxidoreductase and a complete succinate dehydrogenase, as previously described, indicating potential for fumarate respiration. A fermentative, heterotrophic lifestyle has been previously hypothesized, with sugar catabolism genes for glucose and mannose utilization identified in genome QTKG01.1, as well as amino acid utilization via peptidases. Using DRAM, we identified carbohydrate-active enzyme (CAZy) sequences for the utilization of arabinose, chitin, pectin, mixed-linkage glucan and xyloglucan hemicellulose, and polyphenolics in QTKG01.1. Some functional potential is similar to krumholzibacteria_1, in which we identified CAZy sequences for the utilization of chitin, mixed-linkage glucan, alpha-mannan, and polyphenolics. These differences in functional potential between the two genomes can be expected given the degree of taxonomic novelty and low average amino acid identity (40.9%).

## Conclusion

Biogeochemical evidence for methane and iron cycling in Bothnian Sea sediments (23, 26) highlights the importance of microbial processes in controlling greenhouse gas emissions from coastal ecosystems. This investigation of microbial communities from Bothnian Sea sediments under low oxygen concentrations in a bioreactor receiving methane and iron (III) as main substrates revealed functional potential for methane and iron cycling in novel taxa. Moreover, this study brought new hypotheses on the identity and metabolic versatility of microorganisms potentially members of these functional guilds, such as the enriched *Verrucomicrobia, Bacteroidetes* and *Krumholzibacteria*. These results also indicate that the methane biofilter in coastal sediments may be more diverse than previously thought. Finally, our results are compared to recent evidence that bacterial methanotrophs may also be capable of anaerobic oxidation of methane. Future studies, which are needed to elucidate *in-situ* metabolic activity and mechanisms for methane and iron cycling in Bothnian Sea sediments, will benefit from the insights gained in this work.

---

**Figure S1.**
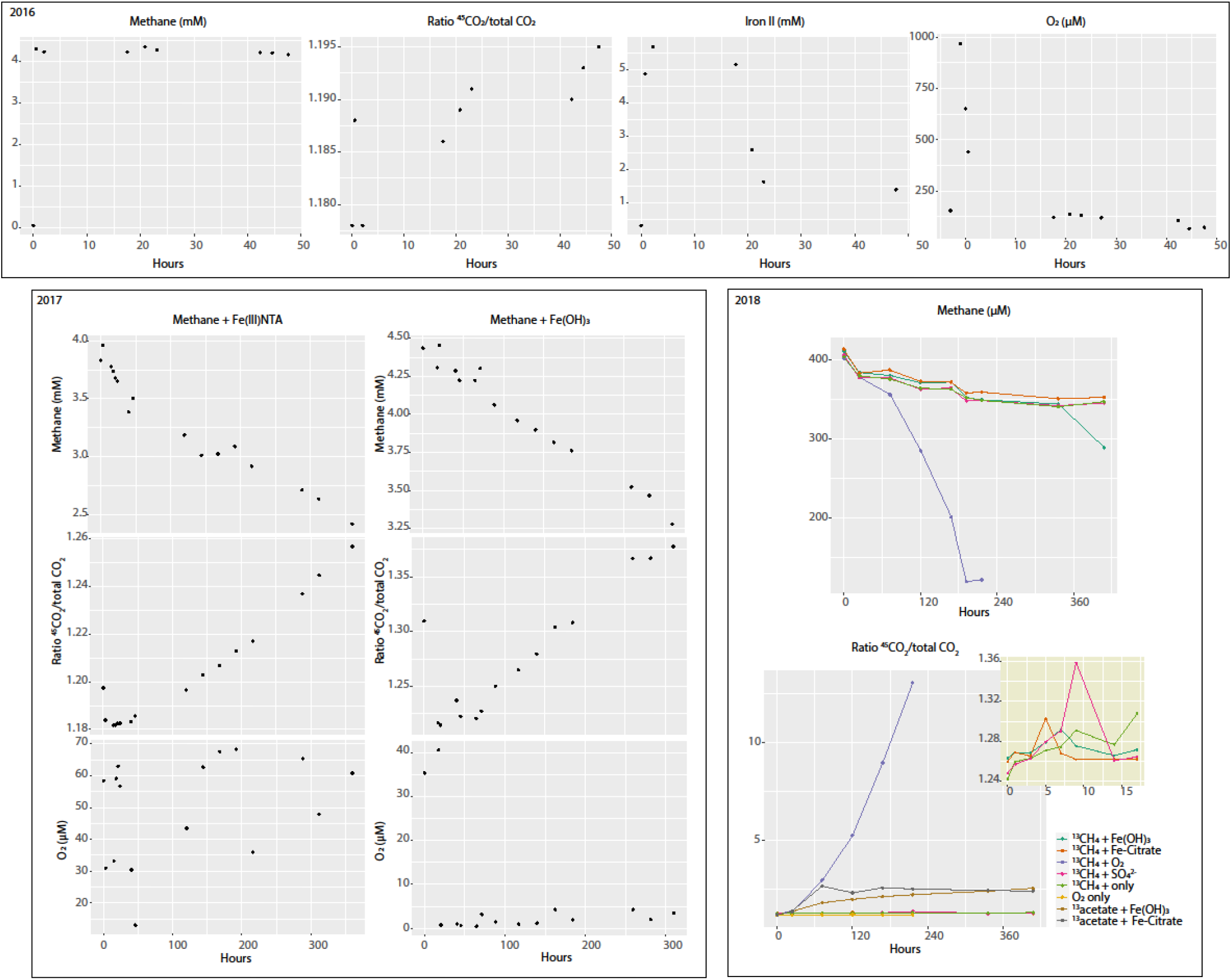
Summary of activity tests. Data are categorized based on the year when measurements were conducted. In 2016 and 2017, whole reactor experiments were performed, in which the reactor was set as a closed system and substrates were followed for several hours. In 2018, bioreactor biomass was incubated into serum bottles with combinations of electron donors and acceptors as indicated in the figure and detailed in the methods section. The inset in yellow displays a zoom in on some incubations as specified in the legend. Oxygen measurements reflect headspace concentrations.

**Figure S2.**
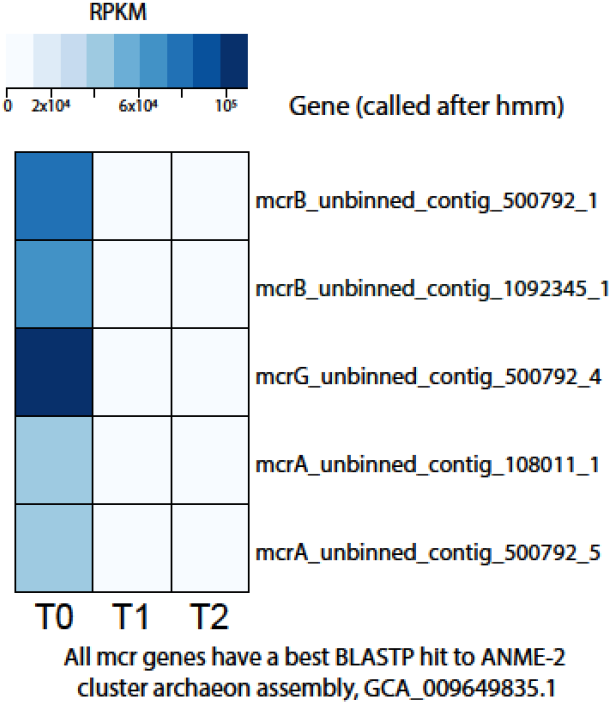
Heat map of *mcr* genes in the three metagenomes (T0, T1, and T2). RPKM values were calculated from read mapping with BBMap and imported into RStudio for heat map construction. Protein sequences were used for blastp against the NCBI RefSeq database.

**Figure S2.**
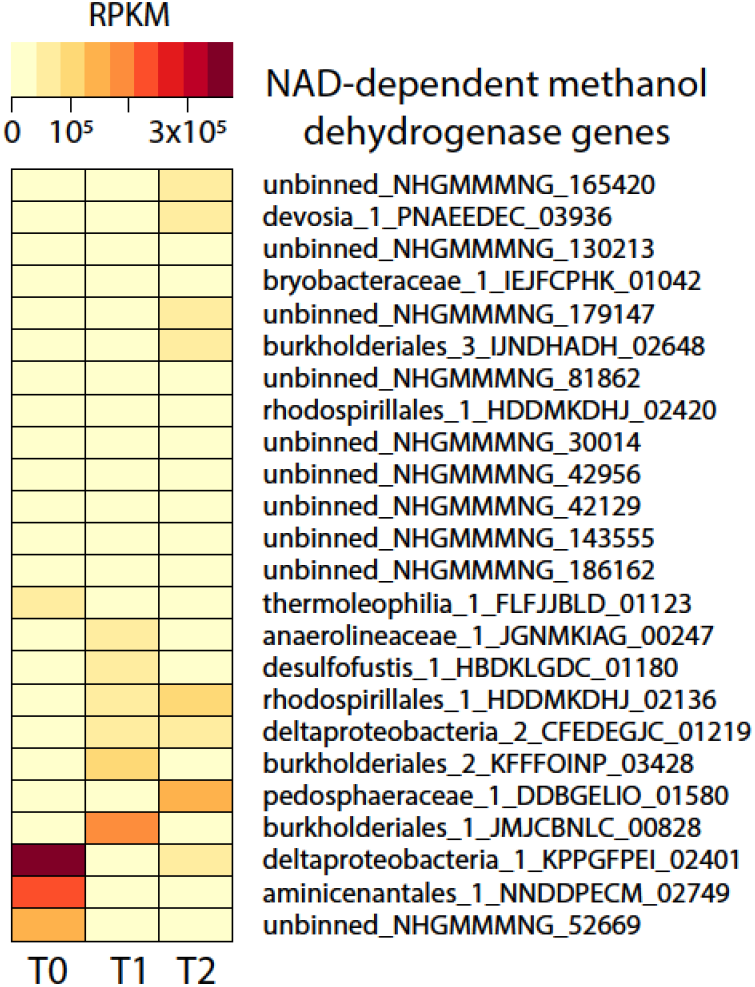
Heat map of NAD-dependent methanol dehydrogenase genes in the three metagenomes (T0, T1, and T2). RPKM values were calculated from read mapping with BBMap and imported into RStudio for heat map construction. Protein sequences were identified via prokka annotations.

## Acknowledgements

We thank Annika Vaksmaa for DNA extractions and useful discussions, Theo van Alen for sequencing the metagenomes, Jeroen Frank for binning support and Matthias Egger for assistance with sampling.

## Author contribution

PDM conducted metagenomic analyses, made figures, deposited data on NCBI, and wrote the manuscript. ADJ inoculated the bioreactor, maintained it for ∼2.5 years and performed activity assays. WKL, NvH and CPS organized the sampling campaign, collected sediments, and informed bioreactor set up and experiments. MSMJ, CUW, and OR planned and guided experiments and analyses. All authors revised the manuscript and agreed on its submission.

